# Conserved Genomic Terminals of SARS-CoV-2 as Co-evolving Functional Elements and Potential Therapeutic Targets

**DOI:** 10.1101/2020.07.06.190207

**Authors:** Agnes. P. Chan, Yongwook Choi, Nicholas J. Schork

## Abstract

To identify features in the genome of the SARS-CoV-2 pathogen responsible for the COVID-19 pandemic that may contribute to its viral replication, host pathogenicity, and vulnerabilities, we investigated how and to what extent the SARS-CoV-2 genome sequence differs from other well-characterized human and animal coronavirus genomes. Our analyses suggest the presence of unique sequence signatures in the 3’-untranslated region (UTR) of betacoronavirus lineage B, which phylogenetically encompasses SARS-CoV-2, SARS-CoV, as well as multiple groups of bat and animal coronaviruses. In addition, we identified genome-wide patterns of variation across different SARS-CoV-2 strains that likely reflect the effects of selection. Finally, we provide evidence for a possible host microRNA-mediated interaction between the 3’-UTR and human microRNA hsa-miR-1307-3p based on predicted, yet extensive, complementary base-pairings and similar interactions involving the Influenza A H1N1 virus. This interaction also suggests a possible survival mechanism, whereby a mutation in the SARS-CoV-2 3’-UTR leads to a weakened host immune response. The potential roles of host microRNAs in SARS-CoV-2 replication and infection, and the exploitation of conserved features in the 3’-UTR as therapeutic targets warrant further investigation.

## INTRODUCTION

The COVID-19 infectious disease outbreak is having a dramatic effect on not only public health, but the global economy as well. The severe acute respiratory distress associated with SARS-CoV-2 infection, the pathogen responsible for COVID-19 illness, was first reported in 2019 (*1*, *2*). As of June 2020, SARS-CoV-2 has been detected in over 188 countries, infecting over 10 million people, and responsible for more than 0.5 million deaths (Johns Hopkins Coronavirus Resource Center, https://coronavirus.jhu.edu/map.html) (*3*). The genome of SARS-CoV-2 is small but complex, encoding structural proteins and regulatory elements whose functions, behavior and interactions with host factors have been studied extensively (*4*, *5*). However, many of these studies have, justifiably, focused on one or another aspect of the SARS-CoV-2 genome, such as the structural proteins it encodes (*6*), its relationships to other viruses (*7*), or its diversity across the locations in which people have been infected (*8*). This leaves room for broader, more integrated approaches to the analysis of the SARS-CoV-2 genome focusing on, e.g., non-coding elements, that could yield insights missed by studies with a singular focus.

The SARS-CoV-2 pathogen is a Coronavirus (CoV), and CoVs are members of the family Coronaviridae. Coronaviridae are divided into four genera based on phylogeny: alphaCoV, betaCoV, gammaCoV, and deltaCoV. CoVs have been detected in a diverse group of hosts from humans, wild mammals (e.g. bats, pangolins, camels, civets) and birds, to farm animals and poultry (*9*, *10*). The betaCoVs are further divided into four lineages: A, B, C, and D. SARS-CoV-2 belongs to betaCoV lineage B and shares moderate genetic similarity with two human pathogenic members, SARS-CoV (lineage B, ~79%) and MERS (lineage C, ~50%), which were responsible for outbreaks of severe respiratory diseases in humans in 2002-2003 and 2012, respectively (*11*). Unlike SARS-CoV-2 infection, human infection by other CoVs causes mild, common cold-like symptoms. For example, the pathogens 229E and NL63, which belong to the alphaCoV, and pathogens OC43 and HKU1, which are within betaCoV lineage A, cause mild symptoms in humans. This suggests that genetic differences between SARS-CoV-2 and related viruses may explain its exceptional infectivity, pathogenicity and elusiveness to effective vaccine and pharmacological mitigation strategies (*12*, *13*).

Many non-coding elements of the SARS-CoV-2 genome have begun to receive attention as potentially informative with respect to the origins and vulnerabilities of the virus. For example, the genomic terminals of CoVs reflect non-coding 5’- and 3’-untranslated regions (UTRs) and encode conserved RNA secondary structures that have unique gene regulatory functions (*14*). The UTRs are shared by both genomic and sub-genomic RNAs and have been suggested to play important roles in viral replication and transcription. The UTRs can also recruit and interact with a range of host and viral protein factors, and may provide long-range RNA-RNA or RNA-protein interactions through circularization of the genome. MicroRNAs (miRNAs) are evolutionarily conserved non-coding RNAs which can repress gene expression post-transcriptionally via partial sequence matches primarily to the 3’-UTR of the target RNAs. In this light, human miRNAs can target viral RNAs and modulate different stages of the viral replication life cycle, positively or negatively (*15*). An example of human miRNA providing a positive influence on viral replication can be found in the hepatitis C virus (HCV), in which the human liver-specific miR-122 stabilizes the 5’-UTR of HCV leading to the promotion of viral replication. Antisense oligonucleotides acting as inhibitors of miR-122 have been developed as antiviral drugs to reduce viral loads in patients (*16*). There are also examples of human miRNAs having the opposite effect. For example, a human miRNA showing a negative influence on viral replication (i.e. positive outcome for the host) has been reported in the Influenza A virus (IAV) H1N1. Five human miRNAs that are highly expressed in respiratory epithelial cells targeting multiple gene segments have been shown to provide inhibitory effects on IAV replication both *in vitro* and *in vivo* (*17*).

We pursued a systematic gene-by-gene comparative analysis, assessing sequence conservation in each region and element of the SARS-CoV-2 genome, including the 5’- and 3’-UTRs. We determined whether each of these regions and elements were broadly conserved across the CoV family, or unique to sub-lineages of CoVs. We also identified mutation hotspots, characterized the likely functional significance of naturally-occurring amino acid substitutions, and assessed evidence for co-evolving mutations across the genome that may impact the stability of the SARS-CoV-2 genome as a whole. Finally, we identified a unique genomic signature residing in an evolutionarily conserved element in the 3’-UTR, which could be involved in host miRNA-mediated interactions and innate immunity response. These findings reveal unique viral and host conserved elements associated with the SARS-CoV-2 genome and warrant further investigation into their possible functional roles during infection as well as potential therapeutic targets.

## RESULTS

### Conserved Sequence Features of the Coronavirus Family

To identify conserved and potentially functional features in the CoV family, *Coronaviridae*, we compared each of the annotated genes and UTR features of the SARS-CoV-2 reference genome (NC_045512.2) against 109 selected CoV family genomes (Table S1). The SARS-CoV-2 reference isolate carries 26 processed peptides and open reading frames (ORFs), as well as 2 UTRs based on NCBI RefSeq annotation. The CoV family genomes we studied were collected from four coronavirus genera (alpha, beta, gamma, and delta) including seven human CoVs (SARS-CoV-2, SARS-CoV, MERS, OC43, HKU1, 229E, and NL63), a number of mammalian CoVs (e.g. bats, pigs, pangolins, ferrets, civets), as well as avian CoVs (e.g. chicken, fowls). The SARS-CoV-2 sequence features were mapped to the CoV family genome sequences through both nucleotide and amino acid sequence alignments using BLAST (*18*), independently of any CoV family genome annotation (Fig. 1).

**Fig. 1.**
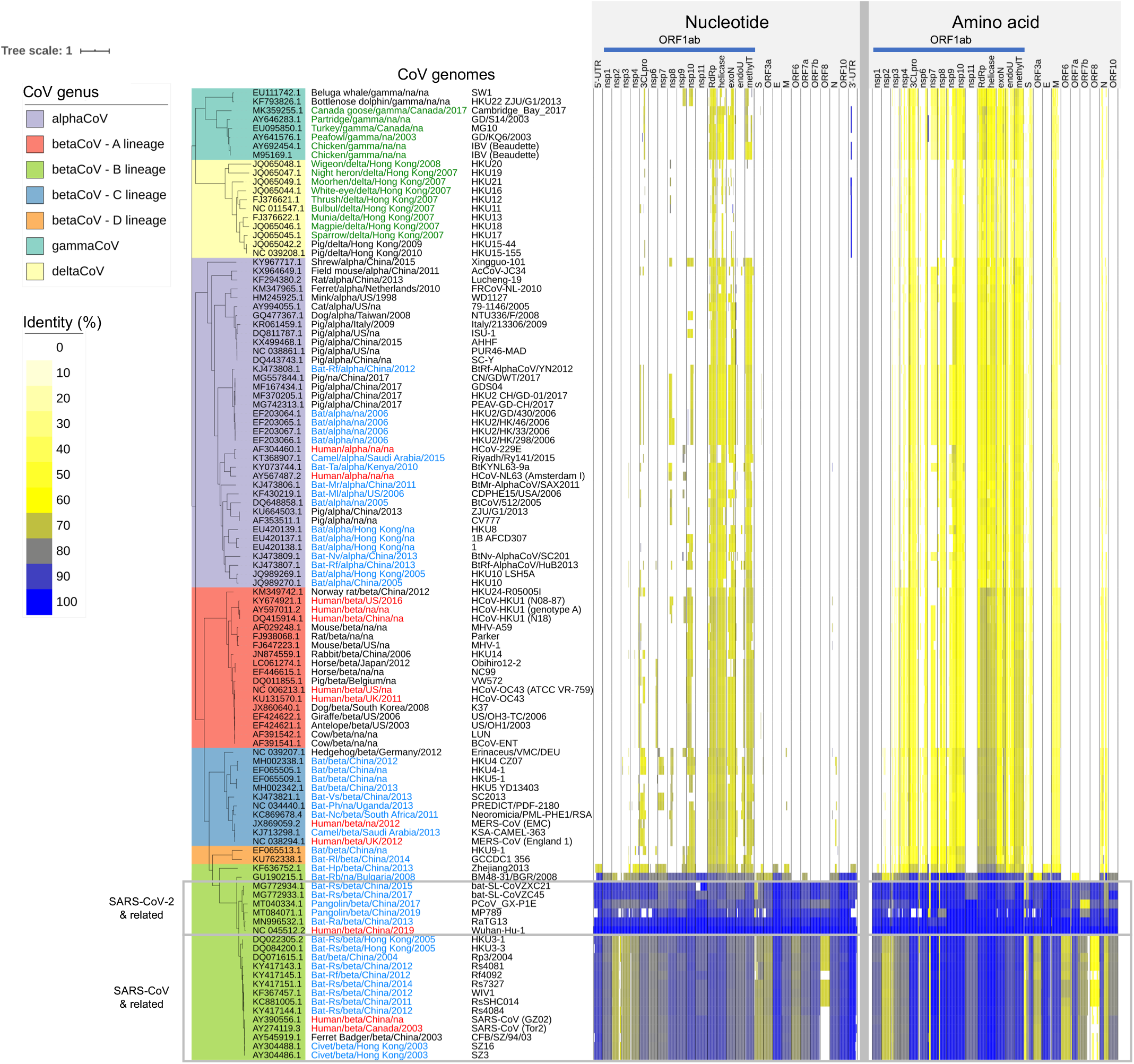
Coronavirus Family Genome Diversity and Conserved Features. The coronavirus family whole genome phylogeny with different genera and sub-lineages represented is provided on the left. Each row corresponds to a different coronavirus family member annotated with host, genus, collection location and year, and the isolate name. The CoV names are color-coded to indicate host species (red: human; blue: bat, civet, camel; green: bird). The columns on the right correspond to gene products and UTR features along the length of the coronavirus genomes with each feature normalized to the same column width. The color intensities indicate the degree of nucleotide and amino acid conservation (i.e. sequence identity) with respect to the SARS-CoV-2 reference genome (NC_045512.2).

The functional element-based conservation analysis results suggested that the 28 total genomic features (i.e., 26 processed peptides and ORFs + 2 UTRs) can be broadly classified into two groups, those that were conserved across all CoV genera (cross-CoV feature group) and those that were conserved only within the betaCoV lineage B (betaCoV lineage B-specific feature group), which includes human SARS-CoV-2 and SARS-CoV, and animal CoVs from bats, pangolins and civets. The cross-CoV feature group showed moderate levels of protein sequence identity across all genera and included nsp3-10, nsp12-16 (RNA-dependent RNA polymerase, helicase, 3’-to-5’ exonuclease, endoRNAse, and 2’-O-ribose methyltransferase), and the structural proteins Spike (S), Membrane (M), and Nucleocapsid (N) (Fig. 1). The betaCoV lineage B-specific feature group mapped uniquely to the betaCoV lineage B, with no sequence similarity detected in other genera at the nucleotide or protein sequence levels. The betaCoV lineage B-specific feature group included non-structural proteins nsp2 and nsp11, accessory proteins ORF3a, ORF6, ORF7a, ORF7b, ORF8, ORF10, the structural Envelope (E) protein, and the 5’- and 3’-UTRs (Fig. 1). Among these, the five most conserved features between SARS-CoV-2 and the betaCoV lineage B isolates in descending order of average nucleotide sequence identity were the 3’-UTR, the E gene, ORF10, the 5’-UTR, and nsp10 at 97.4, 95.1, 93.8, 91.1, and 89.7%, respectively (Table S2). A short stretch (~30 nt) of the SARS-CoV-2 3’-UTR also shared high sequence identities with specific groups of deltaCoVs (from pigs and birds; 97%) and gammaCoVs (from chicken and fowls; 94%) (see next section below). Taken together, these results showed that the nucleotide sequence of both genomic terminals (3’-UTR and 5’-UTR) are exceptionally conserved and unique within the betaCoV lineage B isolates, and therefore suggests they are of functional significance for SARS-CoV-2.

### Notable Signatures in the UTRs of SARS-CoVs and Related Genomes

To investigate the extent of sequence conservation within the genomic terminals of SARS-CoV-2 and related isolates, we performed a multiple sequence alignment (MSA) analysis on 620 near-full-length betaCoV lineage B genomes collected from the NCBI Nucleotide database, which included 361 SARS-CoV-2, 113 SARS-CoV, 75 animal CoVs (e.g. bats, pangolins, civets), and 71 laboratory isolates (Table S1). The 5’-UTR (SARS-CoV-2, 1 to 265 nt) was defined as the 5’-terminal, and both ORF10 and the 3’-UTR together (29558 to 29903 nt) were used for the 3’-terminal analysis. ORF10 was included in the 3’-terminal analysis because ORF10 was a predicted ORF immediately upstream of the 3’-UTR but no ORF10 expression was detected as reported in a comprehensive SARS-CoV-2 transcriptome analysis (*19*). Hereafter, we will refer to the 3’-UTR as a 3’-genomic terminal including both ORF10 and 3’-UTR, and all genomic coordinates will follow the SARS-CoV-2 reference isolate (NC_045512.2) unless otherwise noted.

The MSA analysis of the 3’- and 5’-UTR revealed near-perfect sequence identity of the regions across the betaCoV genomes. Across the nucleotide positions where most genomes (>99%) have sequence alignments (i.e. ignoring positions near both ends of genome where many genomes do not have sequences), 94% of the 3’-UTR positions (234 out of 249) and 84% of the 5’-UTR positions (151 out of 179) shared identical nucleotides amongst 99% of the genomes aligned. Within these conserved regions, a high level of nucleotide diversity was observed at specific positions across the sequence alignments, with 13 and 25 hypervariable positions identified in the 3’- and 5’-UTR, respectively (Fig. 2). These 38 positions altogether showed distinct nucleotide profiles for sub-clades of the betaCoV genomes, and we refer to them as the UTR ‘signatures.’ A total of major 15 UTR signatures and their frequency distribution were determined from the 620 betaCoV genomes (Fig. 2). Based on nucleotide identities, the UTR signatures could be clustered into two distinct groups represented by the SARS-CoV-2 (Wuhan-Hu-1) and SARS-CoV (Tor2) isolates respectively, which harbored 76% non-identical nucleotides (29 out of 38 positions at the UTR signature positions). The UTR signature of the SARS-CoV-2 clade was shared by bat CoV isolates (RaTG13, ZC45, and ZXC21) and pangolin CoV isolates (MP789, GX-P4L, and GX-P1E); and that of the SARS-CoV clade was shared by a different group of bat CoVs (HKU3-1, Rf1, YNLF_31C, and Rs672) (Fig. 2).

**Fig. 2.**
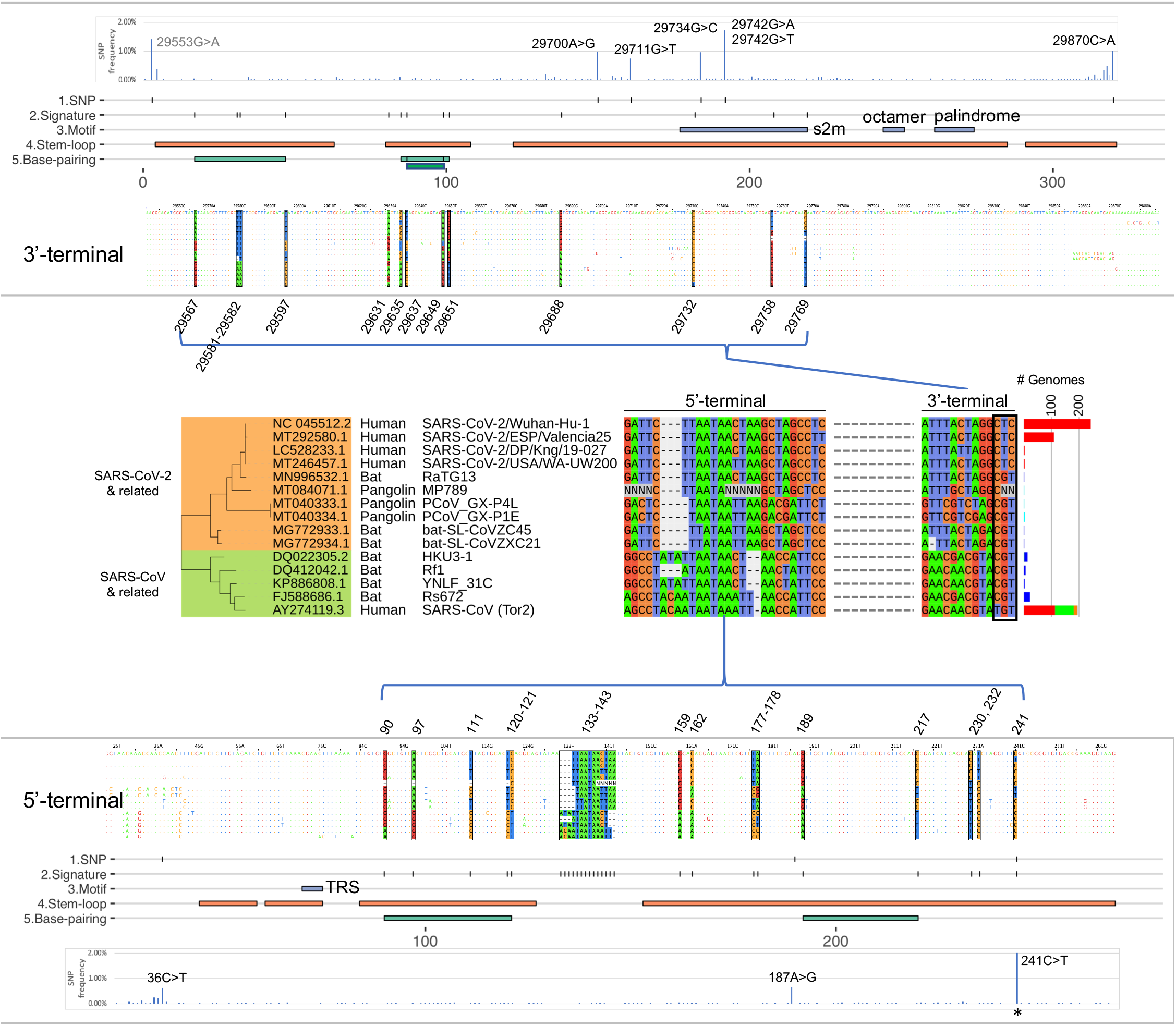
UTR Signatures of BetaCoV Lineage B Genomes. Variant positions in the SARS-CoV-2 5’- and 3’-UTRs and their presence in related SARS-CoV genomes (middle section). Base positions are color coded by the four nucleotides and depicted in their genomic locations for the 3’-UTR (upper panel) and 5’-UTR (lower panel) sequence coordinates. For each panel, the data tracks are: SNV frequency in SARS-CoV-2 genomes based on 18k GISAID genomes analyzed, SNV positions with >0.5% mutation frequency, UTR signature positions, conserved sequence motifs, predicted stem-loops, and predicted complementary base-pairings. The number of betaCoV genomes (# Genomes) carrying each unique signature is shown in a bar plot to the right with the following color codings for host species: red, human; blue, bat; green, laboratory; and orange, civet. The 241C>T SNV is indicated by an asterisk (*) with an observed frequency of 70.2% (outside of the frequency scale shown). The 29553G>A SNV is upstream of the 3’-terminal with no ORF annotation showing moderately high mutation frequency at 1.42%. S2m: Coronavirus 3’ stem-loop II-like motif, TRS: transcription regulatory sequence.

Overlaying the UTR signatures with predicted RNA secondary structures revealed that a majority of the signature positions (71%; 27 out of 38) were located on stem-loop structures, and that 10 positions were involved in complementary base-pairings. Interestingly, we noted that the last three positions (29732, 29758, 29769 nt) of the 3’-UTR signature carried distinct nucleotide combinations for each group of the SARS-CoV-2 (‘CTC’), SARS-CoV (‘TGT’), and the bat CoVs (‘CGT’) isolates (Fig. 2). Notably, these three positions overlapped with a conserved RNA motif S2m (Coronavirus 3' stem-loop II-like motif, Rfam RF00164) previously identified in coronavirus and astrovirus (*20*, *21*). In our analysis, the highly conserved S2m RNA element was also detectable using nucleotide searches among avian and animal CoVs belonging to the gamma and delta genera (Fig. 1). In summary, these results show that the 3’- and 5’-UTRs of SARS-CoV-2, SARS-CoV, and batCoV isolates carry unique signatures involving predicted RNA secondary structures with likely functional and/or regulatory roles.

### UTR Stability and Variant Sites within the SARS-CoV-2 Genome

To investigate SARS-CoV-2 genomic stability, we analyzed genome-wide nucleotide variants amongst isolates collected from the ongoing global outbreak. We performed single nucleotide variant (SNV) discovery by pairwise whole genome alignments using Nucmer on 18,599 whole genome sequences available from the GISAID resource (as of May 29, 2020, https://www.gisaid.org) (Fig. S1, Table S3), and a set of stringent filtering criteria to identify high confidence SNVs (see methods). Variant analysis identified 87 variant (SNV) positions with frequencies >0.5% (or equivalently, occurring in at least 93 genomes). Inspection of the UTR signature positions showed that 37 out of 38 positions were relatively stable within SARS-CoV-2 isolates with variants detected in <0.11% genomes (i.e., 20 isolates or fewer) (Fig. 2). One exception was the variant g.241C>T, which represented one of the signature positions and was originally discovered using 361 SARS-CoV-2 genomes in the betaCoV lineage B analysis above. In our expanded 18k SARS-CoV-2 genome analysis, the variant g.241C>T was detected at a high prevalence of 70.2%. In addition, six variants were identified at five sites in the 3’-UTR (g.29700A>G, g.29711G>T, g.29734G>C, g.29742G>T, g.29742G>A, g.29870C>A) and three in the 5’-UTR (g.36C>T, g.187A>G, g.241C>T) (Fig. 2). Setting g.241C>T aside, the UTR variants were detected at a low frequency, between 0.62 and 1.05%. A very recent paper by Mishra et al. identified two variant positions overlapping with this analysis in the 5’- and 3’-UTRs respectively (i.e. g.241C>T, g.29742G>A/T) (*22*). In this study, all UTR variants were located on predicted stem-loop structures with the exception of g.36C>T in the 5’-UTR. We note that the 29742 position was located within the conserved RNA motif S2m, and carried two alternate alleles, making it a triallelic site (Fig. 2, see Discussion). The alternate allele g.29742G>T was observed with a frequency of 1.05%, and the second alternate allele g.29742G>A at a frequency of 0.67%. Based on whole genome phylogeny analysis, the g.29742G>T and g.29742G>A variants appeared to have arisen in two distinct clades: the g.29742G>T variant was predominantly found in Asia (43% of G>T isolates), and g.29742G>A almost equally split between Asia and North America (40.0 and 39.5% respectively of G>A isolates) (Fig. S2).

The observed SARS-CoV-2 variants were presumably the result of the evolution of the virus and potential selection pressures on those variants during the pandemic given their likely functional impact on some aspect of the behavior of the virus. Imposing a variant frequency threshold of 0.05% or higher (or equivalently, occurring in 10 or more genomes) identified 769 SNVs (Table S4). By considering the number of variant positions per kilobase across gene features, we found that the 3’-UTR, ORF3a, and the 5’-UTR harbored the highest number of variant positions (Fig. 3A). The next group of genes carrying a high SNV density were ORF10, N, and ORF8, which were all immediately upstream of the 3’-UTR. We analyzed two aspects of the 769 SARS-CoV-2 SNVs by classifying them into the types of observed base changes (i.e. A>T, A>G, A>C, etc.), and amino acid consequences (i.e. missense, synonymous, and nonsense) across the SARS-CoV-2 genes and UTRs. By assigning SNVs into different base change categories, we observed a predominance of C>T mutations out of all 12 possible base changes. The C>T mutation bias in SARS-CoV-2 has been previously suggested to be associated with human host RNA-editing activities and the subsequent fixation of the edited nucleotides in the viral RNA genome (*23*). The study by Di Giorgio et al. (*23*) pointed to C>T/G>A and A>G/T>C variants as base modification outcomes of the human APOBEC and ADAR deaminase family activities, respectively. Results from our gene-by-gene analysis confirmed the study’s observations that: (1) C>T variants were the most abundant base change across almost all gene features, and that (2) C>T variants were biased towards the positive-sense RNA strand (Fig. 3B). In other words, C>T variants were more abundant than G>A, which would have been the complementary base changes if C>T variants were to occur in the negative-sense RNA strand. Importantly, our results further revealed that the above two properties did not hold for the 3’-UTR. In the 3’-UTR, we observed that C>T and G>A variants were similarly frequent, and that G>T variants instead were the most dominant base changes followed by the G>A and C>T variants. These results could indicate different selection pressure or regulation of the 3’-UTR from other parts of the genome. In addition, our analysis also detected G>T variants as the second prominent base change type when considering the entire genome. The gene features showing the highest density of G>T mutations were ORF3a, ORF6, N, and 3’-UTR, all of which were located in the last third of the genome. We determined that the average G>T variant density in the last third of the genome (downstream of ORF1ab) was three times higher than the first two-thirds of the genome (entire length of the ORF1ab) (Fig. 3B; Fisher’s Exact Test, p = 2.6e-09). In summary, G>T variants are more enriched towards the 3’-end of the genome.

**Fig. 3.**
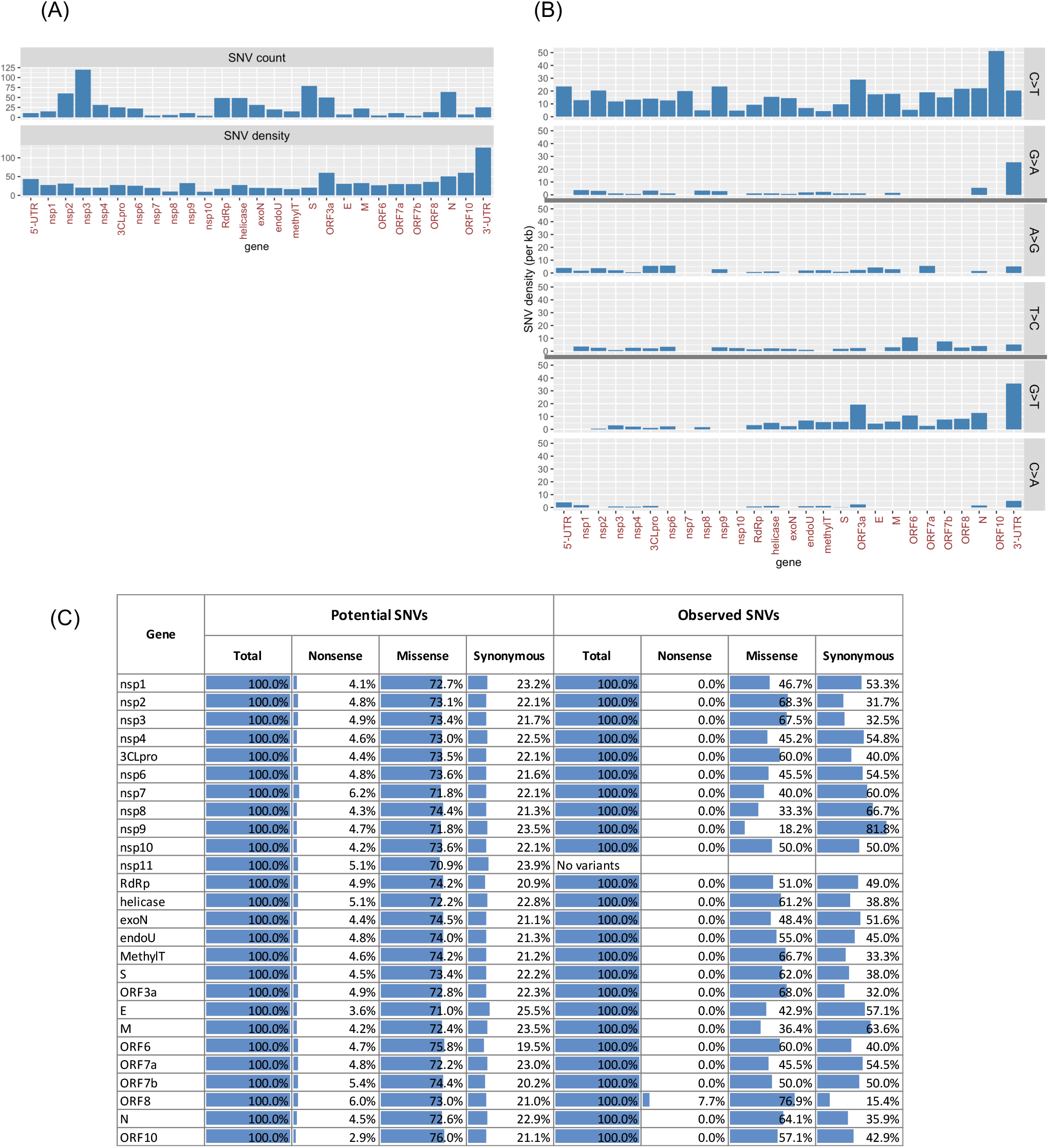
SARS-CoV-2 SNV Properties. A total of 769 SNVs detected at 0.05% mutation frequency of 18K GISAID genomes. (A) SNV counts and density (per kb feature length) across genes and UTRs. (B) SNV density is shown by selected base change types: C>T/G>A, A>G/T>C, and G>T/C>A. A full set of SNV distribution across all 12 base change types is shown in Table S4. (C) Amino acid mutation bias comparing expected (potential) and observed SNVs for each gene or UTR feature.

To investigate whether there are any biases in terms of amino acid (AA) substitutions (i.e. missense, synonymous, and nonsense), we first determined that, if an SNV occurs randomly at any given nucleotide along the genome, the chances that it results in missense, synonymous, and nonsense mutations would be 73, 22, and 5% respectively. We also determined that such a distribution remained the same across all 26 protein-coding gene features (Fig. 3C). By analyzing the observed proportions of AA substitutions of the 769 SNVs, we detected lower than expected nonsense and missense variants across all genes with the exception of ORF8. This result suggested likely purifying selection across the protein-coding genes but not on ORF8. Furthermore, we observed that the deviations of the observed proportions from the expected values varied widely across genes (Fig. 3C). In ORF8, for example, the proportion of missense, synonymous, and nonsense variants were 76.9, 15.4, and 7.7%, respectively, which were similar to expectations. In contrast, for the processed peptide nsp9 (putative function in dimerization and RNA-binding), the corresponding proportions were 18.2, 81.8, and 0%, respectively, revealing fewer missense and nonsense variants than expected. These results suggest that there is likely greatly varying selection and evolutionary pressure on individual SARS-CoV-2 genes. In the nonsense AA setting, only a single nonsense variant out of the 769 SNVs analyzed was detected. The variant was located in ORF8 (p.Q18*). Previous studies have identified multiple variant forms of ORF8 in SARS-CoV and SARS-CoV-related human and animal isolates (*24*), including a 29 nt ORF8 deletion variant that had arisen during the late-phase human transmission of SARS-CoV (*25*). In summary, the characterization of SARS-CoV-2 variants suggests non-random selection pressure and may point to undiscovered driving forces of viral genome evolution originated from the hosts or the virus, and may shed light on the identification of mutations with functional or regulatory roles.

### Analysis of SARS-CoV-2 Variant Combinations

We performed linkage disequilibrium (LD) analysis on SNVs from 18k GISAID genomes using Haploview and identified a total of 34 co-evolving variant groups (referred to as ‘CEV’ groups) with 0.1% or higher genome frequency (Table S3, Fig. 4). Notably, we identified two CEV groups that involved the UTRs as well as other gene features, which may motivate testable hypotheses about functional dependencies or interactions of the associated features. The first CEV group (CEVg1) was 5’-UTR-associated and detected in 69.5% of SARS-CoV-2 genomes, and comprised of four variants that were located in the 5’-UTR (g.241C>T), nsp3 (g.3037C>T, synonymous), the RNA-dependent RNA polymerase (g.14408C>T, p.P323L), and the Spike protein (g.23403A>G, p.D614G) (Fig. 5). In terms of geographic distribution by continent, CEVg1 was predominantly detected in South America (88.2%), Africa (86.8%), Europe (79.6%), North America (66.6%), followed by Oceania (41.6%), and Asia (32.6%) (Fig. S3). CEVg1 was first detected in an isolate collected on February 20, 2020 in Italy (Italy/CDG1/2020; EPI_ISL_412973), and has since showed a dramatic increase from 12.2% to 93.4% between a 3-month period from February to May 2020. An updated analysis on 25k GISAID genomes (as of June 13, 2020) identified an earlier isolate that harbored the same set of variants (England/201040021/202; EPI_ISL_464302) collected on February 3, 2020 in the United Kingdom. The increase of CEVg1 was observed both globally and for each region by continent (Fig. S3). It has been shown that the Spike protein D614G mutation, one of the variants implicated in CEVg1, is able to infect human cells more efficiently and therefore enhances transmission. (*6*). The second CEV group (CEVg5) was 3’-UTR-associated and detected in 0.9% of the genomes, and involved six variants that resided in the leader protein/nsp1 (g.490T>A, p.D75E), nsp3 (g.3177C>T, p.P153L), the exonuclease (g.18736T>C, p.F233L), the Spike protein (g.24034C>T, synonymous), the Membrane protein (g.26729T>C, synonymous), and the 3’-UTR (g.29700A>G) (Fig. 5). CEVg5 was detected in a low proportion of genomes collected in North America (2.4%), Oceania (2.3%), and Europe (0.1%), and not in other regions (Fig. S3). The first such isolate was collected on March 3, 2020 in the US (USA/GA_1320/2020; EPI_ISL_420786). CEVg5 remained as a minor group from March to April, at 1.2, and 0.53%, respectively.

**Fig. 4.**
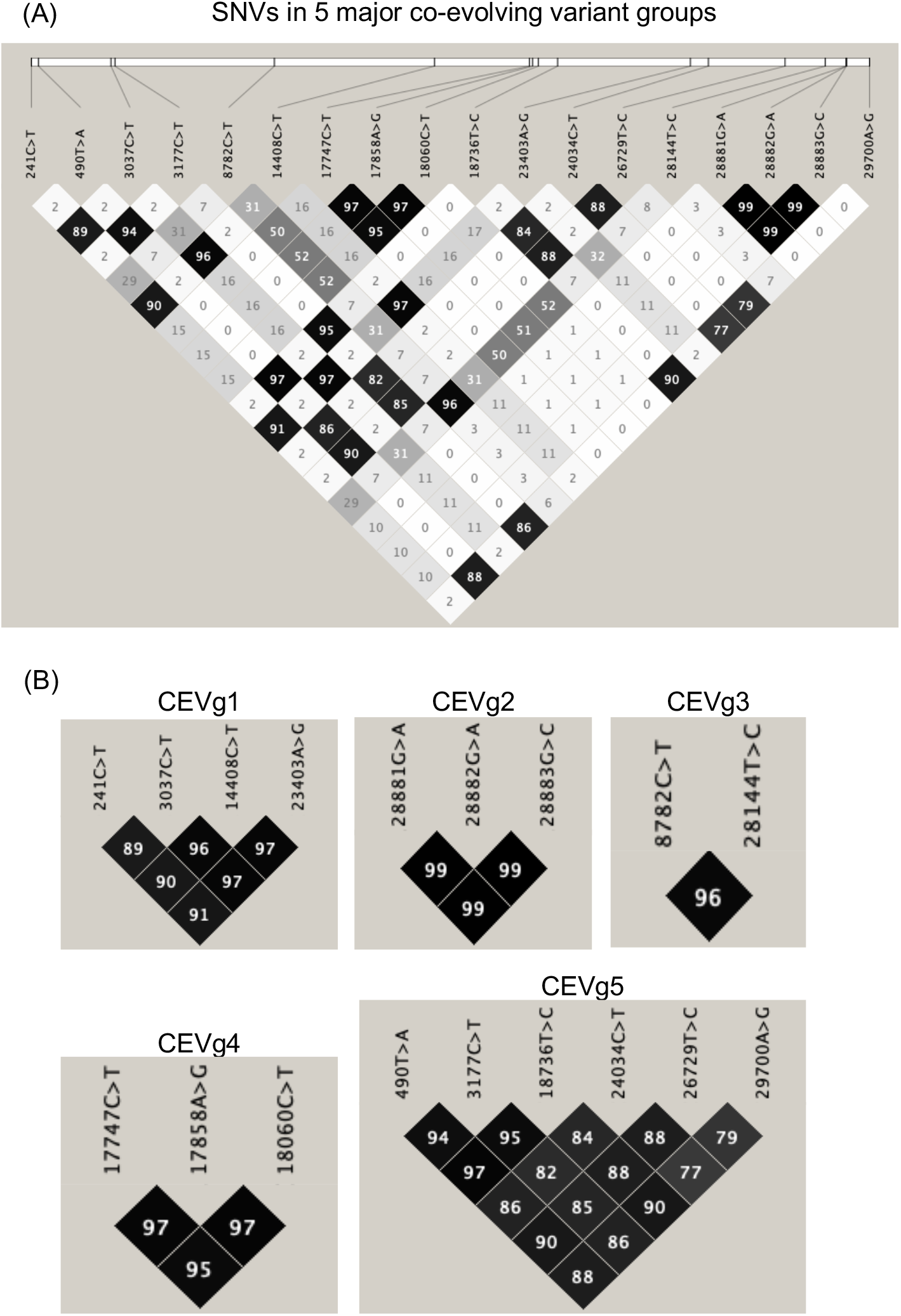
Linkage Disequilibrium (LD) Plots of Co-Evolving Variant Groups. (A) An LD plot of SNVs in the 5 major co-evolving variant groups identified based on 18k GISAID genomes showing the squared coefficient of correlation (r^2^). (B) LD plots of individual co-evolving variant groups.

**Fig. 5.**
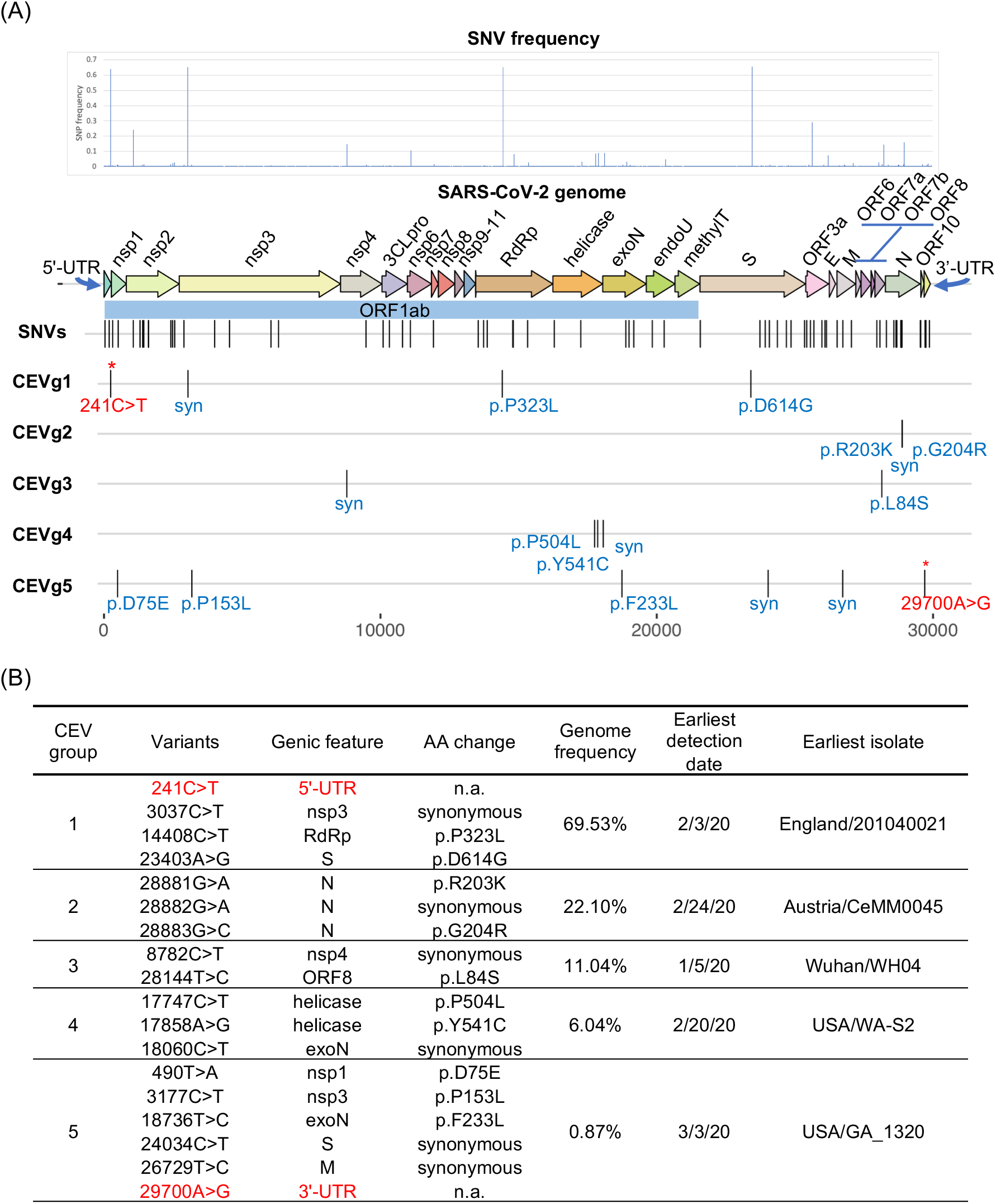
SARS-CoV-2 Co-Evolving and Independent Single Nucleotide Variants (SNVs). (A) SNV frequencies are plotted by the positions in the SARS-CoV-2 genome. The relative positions of five representative co-evolving SNV groups and amino acid consequences are shown. Variant analysis was based on 18,599 genomes from GISAID (May 29, 2020). (B) Five major co-evolving variant (CEV) groups showing different genome frequencies. Two CEV groups involved UTR variants (shown in red font). n.a.: not applicable, syn: synonymous

Three additional CEV groups found in more than 5% of the genomes were identified across gene features (Fig. 5). The first of these three, CEVg2, was detected entirely within the N protein in 22.1% of the genomes. CEVg2 consisted of three consecutive variants g.28881G>A, g.28882G>A, g.28883G>C, which together led to two amino acid substitutions, p.R203K and p.G204R, and the change from one to two positively-charged residues. We predicted the functional impact of the two amino acid substitutions (p.R203_G204delinsKR) using PROVEAN, a variant function prediction tool we previously developed (*26*). The PROVEAN score of −2.856 suggested a deleterious effect as a result of the two amino acid substitutions. These residues were located within a previously identified region (*27*) referred to as the nucleocapsid linker region (LKR, residue 182-247 of SARS-CoV). LKR was identified as a flexible region joining the N- and C-terminal modular regions and included one of three intrinsically disordered regions found in the N protein, and may be involved in phosphorylation, oligomerization, and N to M protein interaction (*27*). Amongst the 18k SARS-CoV-2 genomes, the N protein also harbored the highest number of SNV counts per gene feature (i.e. 12 including co-evolving and single SNVs), of which 8 were found to reside within the LKR. CEVg2 was detected in approximately one-third of the genomes collected in Europe (34.7%) and in South America (28.9%), and was also found between 3.7 to 14.0% in other regions. The first occurrence was detected in an isolate collected on February 24, 2020 in Austria (Austria/CeMM0045/2020; EPI_ISL_437932). CEVg2 has since increased in Europe (Feb to May; 31.9 to 58.9%) and South America (Feb to April; 0 to 36.5%), but has decreased in Asia and Africa (Fig. S3).

The second additional CEV group, CEVg3, included two variants located in nsp4 (g.8782C>T, synonymous) and ORF8 (g.28144T>C, p.L84S), and was found in 11.0% of the genomes (Fig. 5). It has been previously reported by other groups (*28*, *29*). CEVg3 showed different geographic and temporal profiles than those described above. The first occurrence was detected in an isolate collected in Wuhan on Jan 5, 2020 (Wuhan/WH04/2020; EPI_ISL_406801). CEVg3 appeared predominantly in North America (23.7%), Oceania (18.7%), Asia (17.0%), and other regions, and showed a declining trend from 32.3, 13.4, to 1.3%, in January, March and May, respectively (Fig. S3).

The third additional CEV group, CEVg4, consisted of three variants, two in the helicase (g.17747C>T, p.P504L; g.17858A>G, p.Y541C) and one in the exonuclease (g.18060C>T, synonymous), and was detected in 6.0% genomes (Fig. 5). Both amino acid substitutions in the helicase were predicted to be highly deleterious using PROVEAN (p.P504L, score −8.2; p.Y541C, score −8.9). CEVg4 was first identified from an isolate collected on Feb 20, 2020 in the United States (Washington), and most positive genomes (92%, 1036 out of 1124) were detected in North America. The per month occurrence of CEVg4 indicated a decrease between February and April from 8.6, 8.0, to 3.3%, respectively (Fig. S3).

In addition, the nsp2 processed peptide with unknown function carried the highest number of SNV counts (i.e. 10) after Nucleocapsid. A moderately prevalent nsp2 mutation was detected in 22.9% genomes (g.1059C>T, p.T85I), with a predicted deleterious functional outcome (PROVEAN score −4.09) (Table S4). We also noted a deletion of three consecutive nucleotides (g.1605_1607delATG) resulting in an amino acid deletion in nsp2 (p.D268del) was predicted to be deleterious (PROVEAN score −6.370) (Table S4). This deletion of 3 nt, although only identified in a small group of 453 genomes (2.4% global collection), appeared to be highly localized in Europe (95%, 428 out of 453 positive genomes), with only few detected in North America (7 genomes) and Oceania (14 genomes). The deletion was first identified in an isolate collected on Feb 8, 2020 in France (France/RA739/2020; EPI_ISL_410486). A total of 383 genomes were collected from the following regional cluster in proximity: England (124), Netherlands (115), Scotland (102), Northern Ireland (31), and Wales (11). The deletion variant peaked around March in Europe (5.6%) and tapered off in April (2.2%) and May (0.7%) (Fig. S3). In all, our survey of variant positions across 18,599 SARS-CoV-2 genomes suggested that co-evolving and single variants with likely functional impact on viral fitness or pathogenicity were identified across the UTRs and functional elements throughout the genome.

### SARS-CoV-2 UTRs and Human miRNAs as Potential Therapeutic Targets

Viral UTRs and human microRNAs have been explored as therapeutic targets in HCV and other viruses because of their essential roles in viral replication and many additional functional phenomena (*30*). To gain insight into the possible interplay of the SARS-CoV UTRs with host microRNAs in modulating infection pathogenesis, we searched for human miRNAs sharing sequence identity with the UTR sequences of SARS-CoV-2 and SARS-CoV. We used miRBase-specific criteria for BLAST analysis (see methods) for this purpose, and identified from miRbase (*31*) a total of 8 and 7 human microRNAs including sense and antisense matching the 3’- and 5’-UTRs, respectively (Table S5). All except one miRNA-matching region (14 out of 15 miRNAs regions) were located on predicted stem-loop structures (Fig. 6A, B). Sequence matches to the human miRNAs hsa-miR-1307-3p and hsa-miR-1304-3p were located within the broader conserved RNA motif S2m. Interestingly, a recent study of IAV H1N1 provided supporting functional evidence of hsa-miR-1307-3p in mediating antiviral responses and inhibiting viral replication (*32*). We discuss a possible similar role of human miR-1307 in SARS-CoV-2 infection below (see Discussion).

**Fig. 6.**
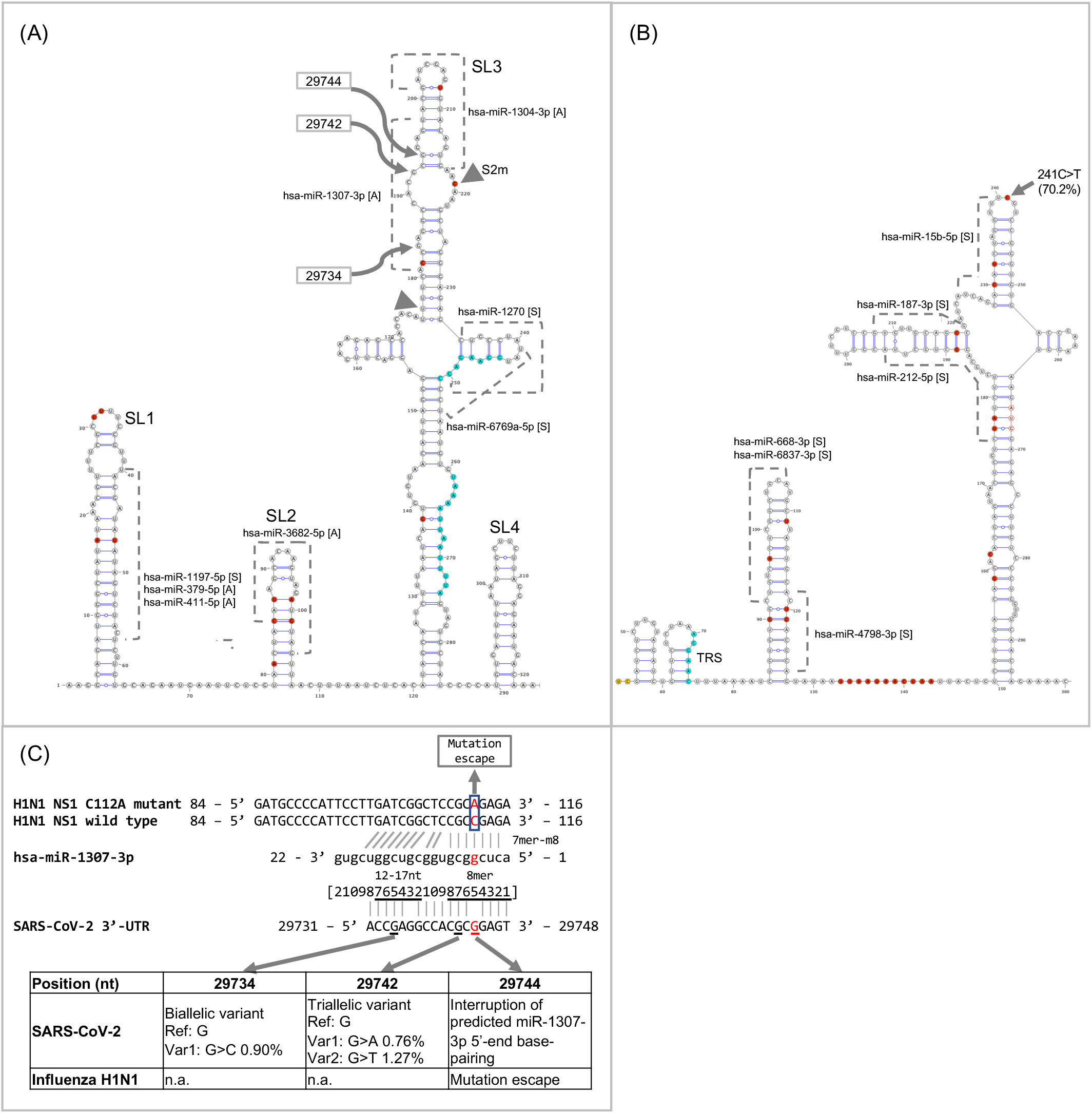
Putative human microRNA interactions with the SARS-CoV-2 UTRs. (A) Predicted secondary structures of the 3’-UTR. (B) Predicted secondary structures of the 5’-UTR. (A, B) Putative human microRNA binding sites with their orientations are shown (A: antisense; S: sense). Nucleotides corresponding to the UTR signatures are colored in red. Sequence features of unknown functions (octamer and palindrome) in the 3’-UTR, and the conserved element TRS in the 5’-UTR are shown in blue. The S2m motif is indicated by two inverted triangles. (C) Putative base-pairings between human miRNA hsa-miR-1307-3p and the SARS-CoV-2 3’-UTR. The base-pairings of miR-1307-3p against the H1N1 NS1 C112A mutant and the H1N1 NS1 wild type sequences were based on (*32*) and updated using bifold prediction (*53*).

We also examined the expression of the 15 identified miRNAs using the human miRNA tissue atlas IMOTA (*33*), which provided categorized miRNA expression levels (i.e. high, medium, low, or not expressed) for 23 human tissues (Table S5). Among the 8 miRNAs with expression data available, three miRNAs (hsa-miR-1307-3p, hsa-miR-1304-3p, and hsa-miR-15b-5p) were reported to be expressed mostly at medium level in all 23 tissues including lung, heart, liver, kidney, and small intestine, some of which were reported to be severely affected during the SARS-CoV-2 infection (*34*, *35*). Furthermore, we searched the miRBase database to determine whether the 15 identified human miRNAs were conserved in other organisms. While 6 miRNAs were not detected in other organisms, 9 miRNAs were found in a number of other mammalian species ranging from 3 to 25 (Table S5). The hsa-miR-1307-3p miRNAs, for example, have been found in 12 other mammalian species in various taxonomic orders such as Primates (e.g. orangutan, chimpanzee, baboon, aye-aye), Artiodactyla (e.g. pig, goat, cow), and others (e.g. bat, dog, rabbit, horse, armadillo). SARS-CoV-2 viral sequences have been detected in dogs from households with confirmed human cases but the dogs remained asymptomatic (*36*). In summary, these results showed that the non-coding UTRs of SARS-CoV-2 harbored sequence similarity that could potentially interact with microRNAs in humans or other species. Further functional assays are needed to delineate whether and how microRNAs are involved in the modulation of viral replication and pathogenesis.

## DISCUSSION

Our SARS-CoV-2 genome-wide analyses demonstrate that ultra-conserved 5’- and 3’-terminal regions of SARS-CoV-2 are shared amongst betaCoV lineage B genomes, including SARS-CoV and different groups of bat CoVs, albeit genome-wide genetic similarity could be as low as ~79%. Notable UTR variant signatures, including complementary-base pairing positions with encoded secondary structures, were identified from representative genomes. The high degree of primary sequence conservation of the UTRs identified in this study, and the predicted RNA secondary structures reported in two recent studies (*37*, *38*), provide strong evidence for conserved functions of the UTRs in the betaCoV lineage B SARS family of viruses. The likely participation of UTRs in long-distance RNA-RNA and/or RNA-protein interactions involving viral and host factors in the replication of CoVs has been proposed and is consistent with our study results and therefore deserves greater attention (*14*).

In addition, our gene-by-gene or functional-element-by-functional-element, comparative analysis of the CoV family provided an account of sequence conservation and dissimilarities in both the nucleotide and amino acid aspects across each functional unit (processed peptides, ORFs, and UTRs) of the SARS-CoV-2 genome. The CoV family reference genomes were collected from multiple sources including NCBI RefSeq (*39*) and previous CoV studies (*40*, *41*), and therefore represent a broad collection of all CoV genera (alpha, beta, gamma, and delta), host species (humans, mammals, and birds), and disease outcomes (human or farm animal outbreaks, or mild symptoms). We believe that our genome-wide sequence analysis is complementary to conventional MSA and phylogenetic analyses (e.g. gene tree) (*4*), or localized window-based analyses (e.g. Simplot) (*2*), which have been used to assess genome/gene sequence conservation. The cross-CoV conservation data generated in this study will provide the basis for a range of follow-up studies, such as determining the functional significance of highly conserved genes and domains (e.g. the E protein), designing vaccine candidates based on protein or RNA conservation, or developing lineage-specific diagnostic markers for community monitoring and interspecies tracing.

Our analyses also suggest that naturally occurring variants in the SARS-CoV-2 genome sequence were relatively low, with approximately 0.3% sites exhibiting variations if one imposes a 0.5% or higher mutation frequency threshold. This is consistent with a low mutation rate of the SARS-CoV-2 RNA-dependent RNA polymerase which likely possesses a proof-reading function similar to SARS-CoV (*42*). The observation that the SARS-CoV-2 UTRs harbored a higher frequency of natural variations (3’-UTR, 2.6%; 5’-UTR, 1.2%) when compared to the overall genome-wide mutation rate of 0.3%, was likely due to lower evolutionary constraints present in the non-coding UTRs than genes in the protein-coding regions. The recent report suggesting the influence of human RNA-editing activities on viral genome mutations has provided some explanations for the overall mutation biases observed (i.e. C>T predominance) (*23*).

Identifying possible therapeutic targets in non-coding regions of a genome has been pursued with other RNA viruses (*30*), and our investigations suggest possible SARS-CoV-2 UTR interactions with human miRNAs. We used a bioinformatics approach to identify genomic regions sharing strong sequence identity (>=18 nt) to human miRNAs as represented in miRBase (*30*, *31*). Because the mature miRNAs can recognize and bind to a target RNA site through canonical or non-canonical matching positions, our initial analyses used sequence identity as an all-inclusive guiding parameter for the human miRNA screen. More specialized miRNA target prediction programs and analyses (e.g. TargetScan, miRanda) should be applied in follow-up studies.

We identified a putative hsa-miR-1307-3p binding site in the 3’-UTR of SARS-CoV-2 with strong sequence identity that exhibits 16 nt of Watson-Crick base-pairings out of the first 18 nt of the miRNA (Fig. 6C). The putative binding site spanned a conserved RNA motif S2m, which was also found in the 3’-UTR of subsets of betaCoVs (e.g. SARS-CoV), gammaCoVs (e.g. Infectious Bronchitis Virus from chicken), and deltaCoVs (e.g. birds, pigs). The S2m motif had been previously identified as a conserved element in other CoVs and astrovirus (*20*, *21*). For some of the CoV genomes, due to a lack of high quality sequences available from the genomic terminals (i.e. non-ambiguous bases), the actual frequency or taxonomic distribution of the S2m and other conserved RNA elements present in the UTRs may have been underestimated. Ongoing efforts to collect and whole-genome sequence the repertoire of naturally-occuring CoV isolates from wild animals including bats (*43*) should help to shed new lights into the evolution of CoV functional elements.

Previous studies have associated hsa-miR-1307-3p miRNA with cancer progression as well as lung function. miR-1307 was originally discovered as a novel human miRNA up-regulated in Epstein-Barr virus (EBV)-positive nasopharyngeal carcinomas (*44*), and also suggested to be associated with the progression of prostate cancer (*45*). mir-1307 expression has been shown to be dysregulated in newborns with chronic lung disease (*46*). Importantly, the study by Bavagnoli et al. demonstrated a functional role of miR-1307 in the regulation of viral replication in the influenza A virus H1N1, which was the pathogen responsible for the 2009 H1N1 pandemic (*32*). Their study predicted sequence complementarity of miR-1307 to the H1N1 nonstructural protein 1 (NS1), which functions to limit interferon and proinflammatory responses thus allowing the virus to evade host innate and adaptive immunity and replicate efficiently in infected cells. The same study also showed that miR-1307 overexpression had regulatory effects on both the virus and host cells. First, miR-1307 overexpression could reduce NS1 expression and inhibit wild type H1N1 replication, but had no effects on the NS1 C112A mutant which carried a nucleotide mismatch to the 5’-region of miR-1307 (Fig. 6C). Second, the overexpression of miR-1307 (in a stably transfected lung cell line) could induce genes involved in cell proliferation, apoptosis, and the regulation of inflammatory and interferon responses. Taken together, the study concluded that the C112A variant was a viral escape mutation for miR-1307 regulation. Furthermore, the study reported that the C112A mutant was significantly associated with the severe clinical symptom acute respiratory distress syndrome, and represented close to one-third of influenza strains that primarily circulated locally in Northern Italy during the 2010-11 influenza season.

In SARS-CoV-2, it is notable that an interruption of base-pairings at 29744 nt to the 5th position of miR-1307 sequence coincided with the location of the C112A mutation in H1N1 (Fig. 6C). It could be hypothesized that SARS-CoV-2 shared a common host defense mechanism as in H1N1 which was mediated by host cellular miRNA regulation, and that SARS-CoV-2 carried an allele having weakened regulation by the human miR-1307 because of the nucleotide mismatch. In support of this hypothesis, our population analysis of SARS-CoV-2 variations identified two nearby mutations at positions 29742 and 29734 nt, which corresponded to the 7th and 15th positions of miR-1307, respectively. Mutations that occurred at these two sites could presumably further disrupt the hypothesized base-pairings with miR-1307 to escape from binding and inhibition. So far, the mutations were detected at a low frequency (0.76-1.3%) in the ongoing outbreak. In all, whether SARS-CoV-2 and H1N1 infections share a similar host defense mechanism mediated by host miRNA regulations, or human population variations of hsa-miR-1307-3p could be associated with the severity of clinical symptoms, are presently not known and warrant further investigation.

## MATERIALS AND METHODS

### Coronavirus family sequence conservation analysis

The SARS-CoV-2 NCBI RefSeq genome (NC_045512.2) was used as the reference. For gene-by-gene analysis, each of 28 annotated genomic features (ORFs, processed peptides, and UTRs) of SARS-CoV-2 was searched against the 109 representative CoV genomes collected from four genera (alpha, beta, gamma, and delta) (Table S1) using NCBI BLAST+ (blastn and tblastx; v2.9.0) with an E-value threshold of 1e-3. The MSA of the 109 CoV family genome sequences was performed using Clustal Omega (v1.2.4) (*47*). The maximum likelihood phylogeny tree was constructed using RAxML (v8.2.11) with 100 bootstraps under the GTRGAMMA model (*48*). The tree was visualized using iTOL (*49*).

### SARS-CoV-2 genomic terminal sequences

In the context of this study, the 5’-terminal (1 to 265 nt) corresponded to the annotated 5’-UTR. The 3’-terminal (29,558 to 29,903 nt), which was also denoted as 3’-UTR, corresponded to the annotated ORF10 and 3’-UTR of the SARS-CoV-2 reference genome (NC_045512.2).

### Collection of betaCoV lineage B genomes and UTR analysis

A total of 693 betaCoV genome sequences were initially collected from the NCBI Nucleotide database (nt database, as of April 15, 2020). Genome sequences were collected using the entire SARS-CoV-2 genome sequence as the query for blastn search and requiring that most of the query sequence length and both UTR regions were aligned sufficiently for sequence comparison (i.e. at least 85% of query sequence is covered; an alignment starting from 130 or smaller nt position exists; and an alignment ending at 29700 nt or higher nt position exists). An MSA was performed on the collected 693 genome sequences including SARS-CoV-2 reference genome using Clustal Omega (v1.2.4). For the 3’- and 5’-UTR regions, variable positions were defined as any positions where 5% or more genomes showed nucleotide differences from the reference (excluding ambiguous nucleotides such as Ns). Positions near either end of the genome (i.e. <87 nt or >29806 nt) were excluded since over 1% of the genomes do not have aligned sequences and therefore the MSA may not be of high quality. Finally, after filtering out the genomes having ambiguous nucleotides on the defined variable positions on UTRs, 620 genomes were used as the final genome set for UTR signature analysis. Note that a pangolin CoV (MT084071.1) was included albeit having ambiguous nucleotides because it appeared to be one of likely close relatives of SARS-CoV-2 and also carried a unique UTR signature.

### Prediction of UTR secondary structure

RNA secondary structure prediction was performed using the RNAfold web server (http://rna.tbi.univie.ac.at/cgi-bin/RNAWebSuite/RNAfold.cgi) with the default basic option to calculate “minimum free energy (MFE) and partition function”. The predicted SARS-CoV-2 5’- and 3’-UTR structures previously reported by (*4*) were used to adjust the prediction.

### SARS-CoV-2 variant analysis

A total of 34,217 SARS-CoV-2 genome sequences and their associated metadata were obtained from the GISAID (https://www.gisaid.org/) on May 29, 2020. A data sanitization and filtering step was performed which included: removing gaps (dash and space characters), filtering out genomes from non-human host, and keeping only high-quality genomes (i.e. requiring a genome to be longer than 29kb, and containing less than 1% Ns and no other ambiguous nucleotides such as B and W). Each of the remaining 18,599 high-quality genomes was aligned with the reference genome to identify variants using the nucmer and show-snps functions of the MUMmer package (v3.23) (*50*). Sequence variants identified within the poly-A tail or near either end of sequence (within 10 nt from either end) were ignored. In addition, an MSA of the 18,599 genomes was built using MAFFT (v6.861b), which was used for independent validations of major mutation positions (*51*). For each sequence variant, the mutation effects on gene products (i.e. genic location and amino acid change if applicable) was analyzed using in-house scripts. The functional impact of amino acid substitutions and indels were predicted using PROVEAN (*26*). Linkage disequilibrium (LD) analysis was performed to identify co-evolving variants among SNVs with frequency of 0.1% or higher using Tagger implemented in Haploview (v4.2) (*52*). Non-biallelic sites needed to be excluded from the LD analysis, and a set of 140 genomes with rare mutations on the major mutable sites, causing the sites to become non-biallelic, were also excluded.

### Protein-coding SNV Analysis

Each of the identified protein-coding SNVs was analyzed to determine its amino acid consequence (missense/synonymous/nonsense) using in-house scripts. For the estimation of amino acid consequences under the assumption of random mutations (i.e. to enumerate all potential SNVs given the sequence context of the SARS-CoV-2 genome), all 3 possible SNVs on every nucleotide position on all coding sequences from the start codon to the last codon before stop codon were included in the analysisd.

### Identification of putatively interacting human microRNAs

The UTR sequences of SARS-CoV-2 and SARS-CoV were used to search against the miRBase mature RNA sequences (Release 22.1) (*31*) using blastn with the following parameters set for short sequences: “-penalty −4 -reward 5 -gapopen 25 -gapextend 10 -dust no -soft_masking false.” For cross-species conservation analysis in other organisms, we searched the miRBase database requiring 18 or more bases matched with 100% sequence identity.

### Statistical Analysis

To test for the significance of the G>T mutation bias toward the 3’-end of the genome, the proportion of G>T mutations out of summed gene lengths was compared between ORF1ab (60 mutations out of 21,326 nt) and the remaining ORFs (66 mutations out of 7,974 nt) using the Fisher’s exact test implemented in fisher.test() function in the R stats package (v3.6.1).

## Definitions of all symbols, abbreviations, and acronyms

AA: Amino acid
CoV: Coronavirus
CEVg: Co-evolving variant group
E: Envelope protein
EndoU: EndoRNAse
ExoN: 3’-to-5’ exonuclease
HCV: Hepatitis C virus
IAV: Influenza A virus
LD: Linkage disequilibrium
M: Membrane protein
miRNA: microRNA
MethylT: 2’-O-ribose methyltransferase
MSA: Multiple sequence alignment
N: Nucleocapsid protein
nt: Nucleotide
ORF: Open reading frame
RdRp: RNA-dependent RNA polymerase
S: Spike protein
SNV: Single nucleotide variant
UTR: Untranslated region

## General

We thank all researchers who have contributed SARS-CoV-2 genome sequences to the GISAID database (https://www.gisaid.org).

## Author contributions

NJS conceived of the study; NJS and AC contributed to overall study design; AC, YC, and NJS identified the appropriate analytical methods and resources; AC and YC conducted the analyses; AC, YC, and NJS interpreted the results of the analyses; and AC, YC and NJS wrote the paper.

## Data and materials availability

Genome sequence data are available through NCBI and GISAID.

## SUPPLEMENTARY MATERIALS

**Fig. S1.**
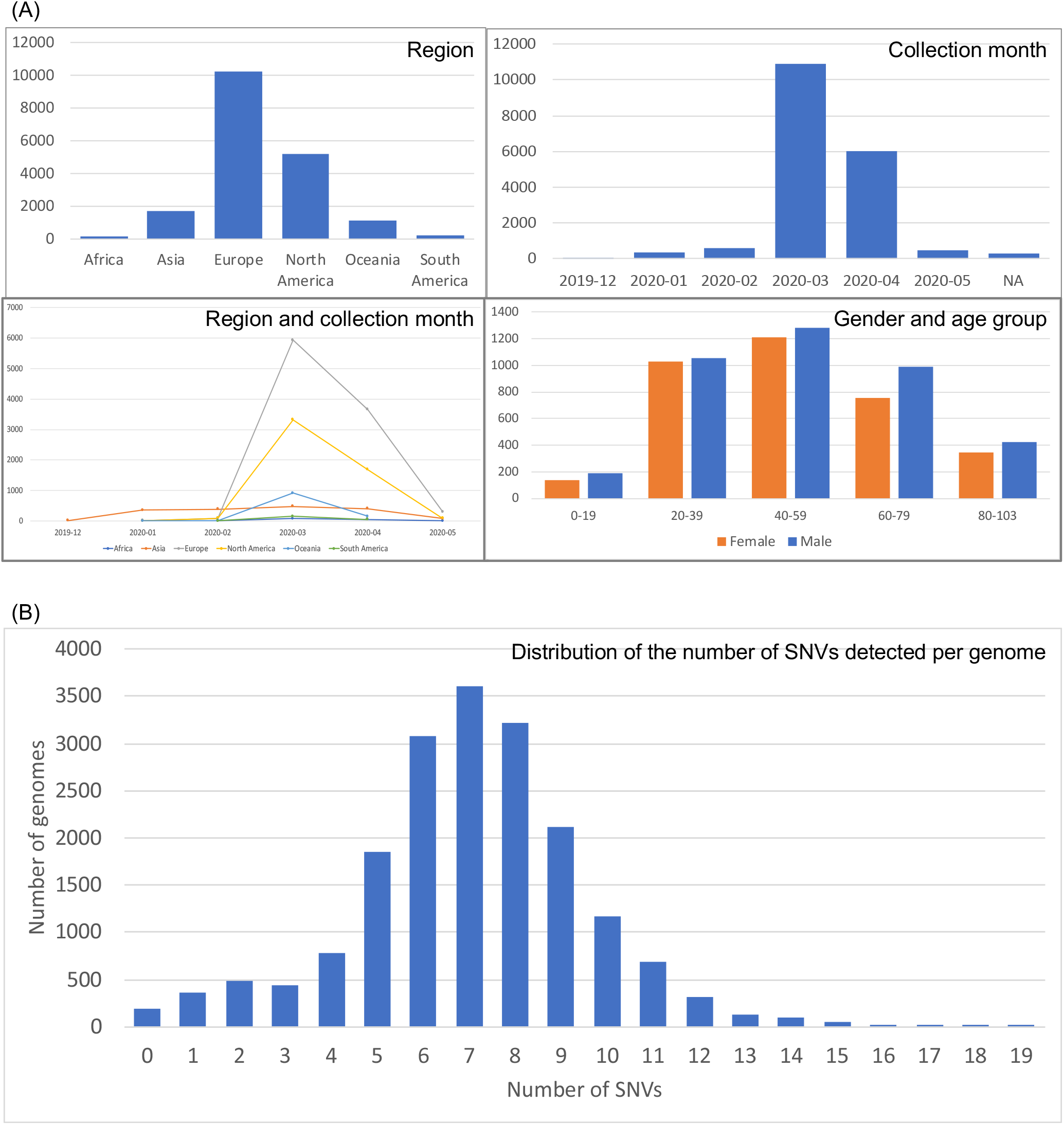
Summary of the 18,599 GISAID SARS-CoV-2 genomes analyzed. (A) Genome distribution by region, collection month, gender, and age group. (B) Distribution of the number of SNVs detected per genome, only including variants detected in two or more genomes (i.e. excluding SNVs unique to a single genome).

**Fig. S2.**
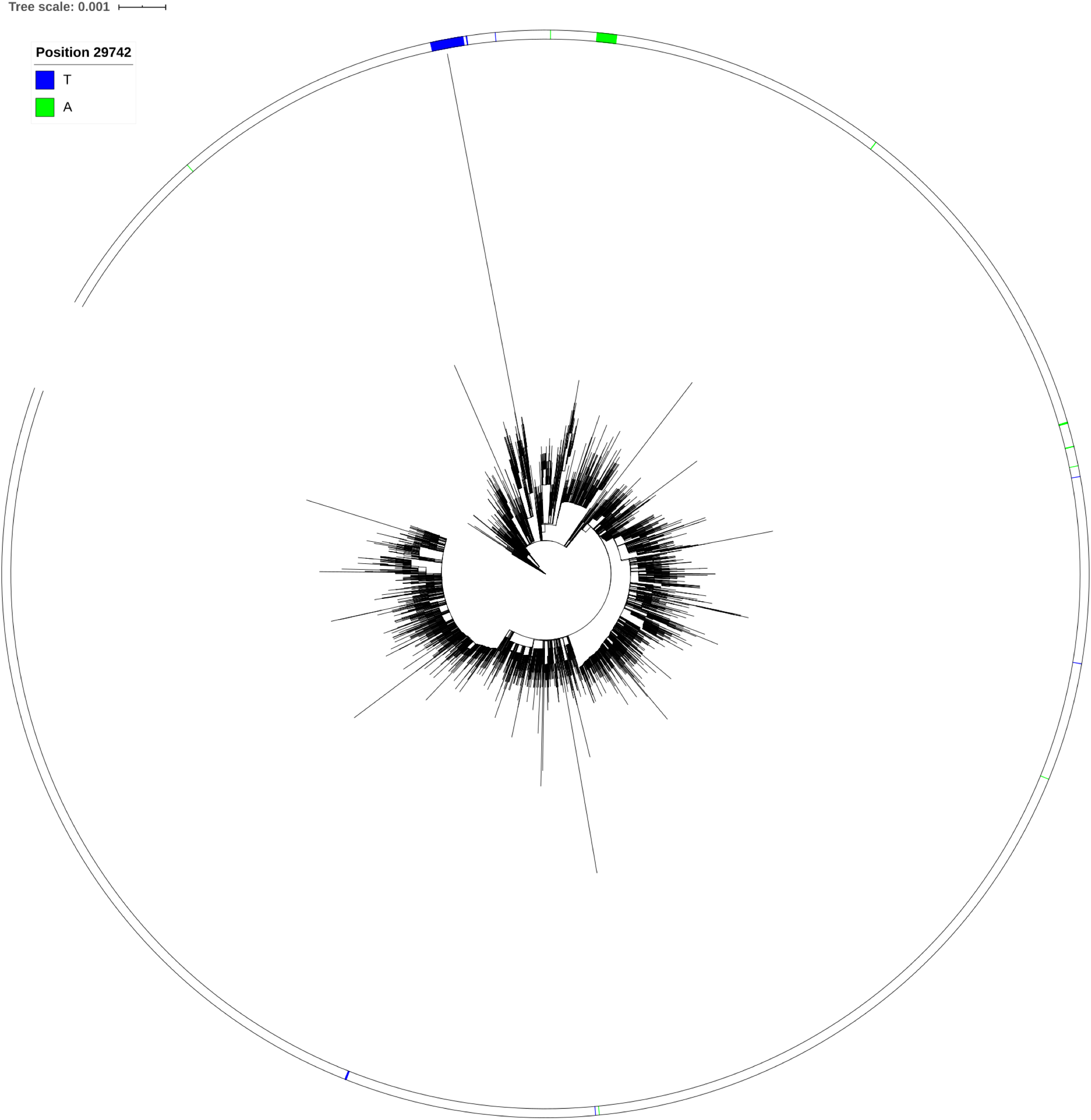
Whole genome phylogeny analysis of SARS-CoV-2 genomes. A maximum likelihood phylogeny tree was constructed from 18k GISAID genomes using RAxML. Genomes carrying the g.29742G>T (blue) and g.29742G>A (green) variants are shown.

**Fig. S3.**
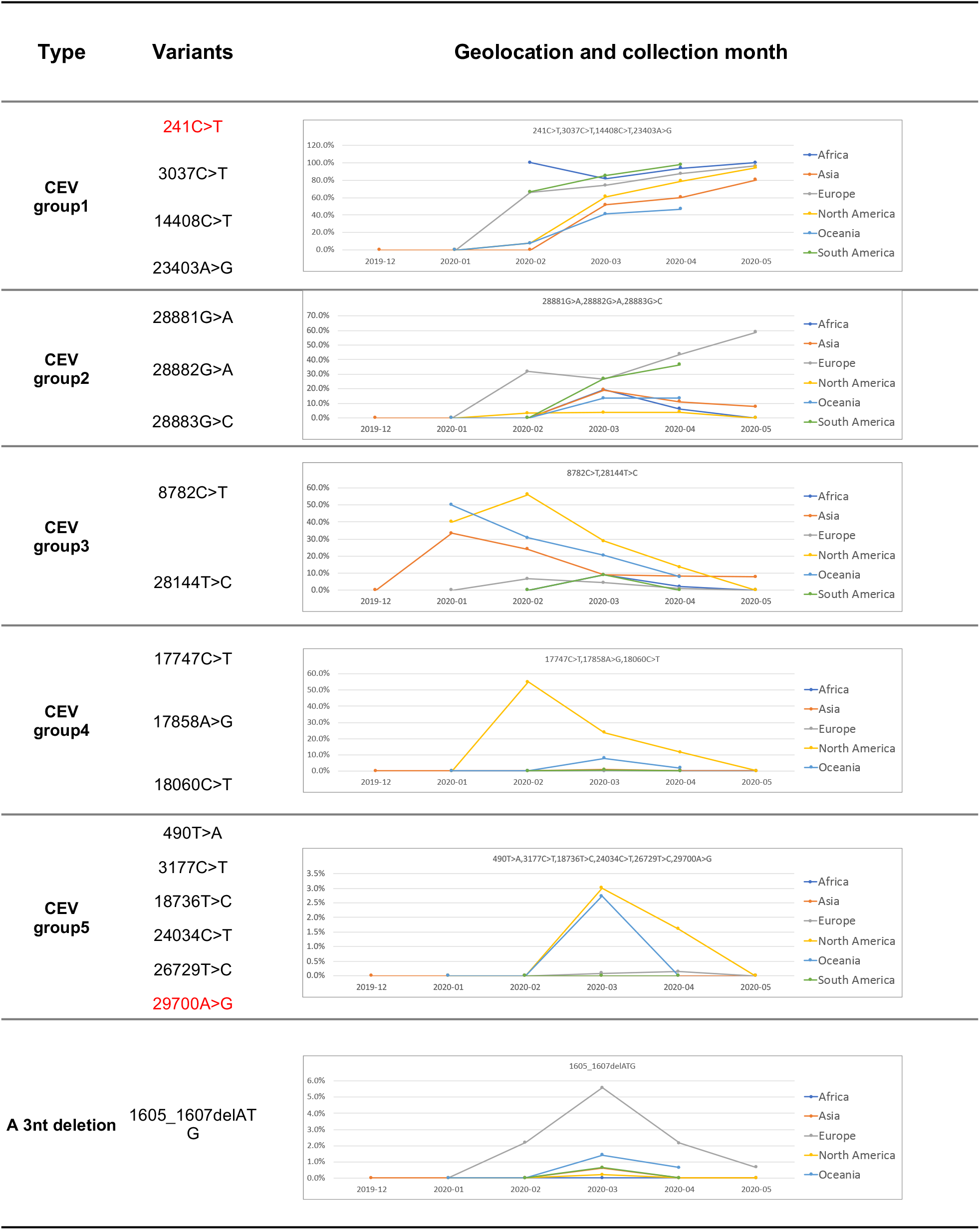
Geographical and temporal distribution of genomes harboring SARS-CoV-2 co-evolving variants. Genome distribution is shown for five co-evolving variant groups and a 3 nt deletion.

**Table S1. CoV genomes collected from NCBI**. (A) 109 reference CoV genomes. (B) 620 betaCoVs genomes.

**Table S2. Sequence comparison of coronavirus family genomes to SARS-CoV-2**. A SARS-CoV-2 gene-by-gene BLAST analysis was performed against the CoV family genomes at nucleotide and amino acid sequence levels. (A) Nucleotide sequence identities. (B) Nucleotide sequence length coverages. (C) Amino acid sequence identities. (D) Amino acid sequence length coverages.

**Table S3. SARS-CoV-2 genomes and metadata obtained from GISAID**. The list included 18,599 genomes analyzed in this study.

**Table S4. SARS-CoV-2 mutations**. The list shows the mutation frequency determined in 18k GISAID genomes, and descriptors for gene locations, amino acid consequences, the earliest detected isolate, and genome counts by exposure region, sex, age group, and collection month. (A) SNVs with 0.05% or higher mutation frequency. (B) Co-evolving variant groups among SNVs with 0.1% or higher mutation frequency. (C) Indels with 0.5% or higher mutation frequency. (D) SNV counts and density per genomic features for all 12 possible base change types.

**Table S5. Putative human miRNAs showing sequence similarities with SARS-CoV or SARS-CoV-2 UTRs**. (A) A list of putative human miRNAs. (B) Human miRNA tissue expression levels collected from the IMOTA database. (C) Human miRNA cross-species conservation analysis.

